# Insights into The Codon Usage Bias of 13 Severe Acute Respiratory Syndrome Coronavirus 2 (SARS-CoV-2) Isolates from Different Geo-locations

**DOI:** 10.1101/2020.04.01.019463

**Authors:** Saif M. Khodary, Ali Mostafa Anwar

## Abstract

Severe acute respiratory syndrome coronavirus 2 (SARS-CoV-2) is the causative agent of Coronavirus disease 2019 (COVID-19) which is an infectious disease that spread throughout the world and was declared as a pandemic by the World Health Organization (WHO). In this study, we performed a genome-wide analysis on the codon usage bias (CUB) of 13 SARS-CoV-2 isolates from different geo-locations (countries) in an attempt to characterize it, unravel the main force shaping its pattern, and understand its adaptation to *Homo sapiens*. Overall results revealed that, SARS-CoV-2 codon usage is slightly biased similarly to other RNA viruses. Nucleotide and dinucleotide compositions displayed a bias toward A/U content in all codon positions and CpU-ended codons preference, respectively. Eight common putative preferred codons were identified, and all of them were A/U-ended (U-ended: 7, A-ended: 1). In addition, natural selection was found to be the main force structuring the codon usage pattern of SARS-CoV-2. However, mutation pressure and other factors such as compositional constraints and hydrophobicity had an undeniable contribution. Two adaptation indices were utilized and indicated that SARS-CoV-2 is moderately adapted to *Homo sapiens* compared to other human viruses. The outcome of this study may help in understanding the underlying factors involved in the evolution of SARS-CoV-2 and may aid in vaccine design strategies.

## 1. Introduction

Baltimore classified viruses into six classes by the means of their genomes. One class is shared between RNA and DNA viruses, while three of them are occupied solely by RNA viruses reflecting their great diversity and different replicative mechanisms (Baltimore, 1971). Over the past few decades, many human infectious diseases including ebola virus disease (EVD), avian influenza, severe acute respiratory syndrome (SARS) resulted from the interspecies transmission of zoonotic RNA viruses (Epstein et al., 2005; Leroy et al., 2005; Subbarao et al., 1998). Most recently, the new pandemic (COVID-19) caused by severe acute respiratory syndrome coronavirus 2 (SARS-CoV-2) has emerged in Wuhan, China in December 2019 and spread to 199 other countries, areas or territories with 462,684 confirmed cases of infection and 20,834 confirmed deaths globally up today (https://www.who.int/emergencies/diseases/novel-coronavirus-2019, 26 March 2020, 20:33 EET). Preliminary phylogenetic analysis showed that SARS-CoV-2 most closely related viruses were (bat-SL-CoVZC45) and a SARS-like beta-coronavirus of bat origin (bat-SL-CoVZXC21). Many encoded proteins revealed a high sequence identity except for the spike (S) protein and protein 13 (80% and 73%, respectively) between SARS-CoV-2 and other bat-derived coronaviruses (Lu et al., 2020). Two probable scenarios are suggested to explain the origin of SARS-CoV-2, natural selection in an animal host either before or following a zoonotic transfer (Andersen et al., 2020).

Coronaviruses (CoVs) are a family of enveloped, single-stranded RNA viruses with the largest genomes (~30 kilobases in length) among other RNA viruses. They are known to cause infection in many avian and mammalian hosts, including humans (Weiss and Navas-Martin, 2005). CoVs genome is organized in 10 open reading frames (ORFs). Four of them (S, M, E, and N) encode for structural proteins: namely, the spike (S) protein, membrane (M) protein, envelope (E) protein and the nucleocapsid (N) protein. The (S) protein has two functions; attachment to the receptors of host cells, and activating the fusion of the virion membrane with host cell membranes (Cavanagh, 2005). The (M) protein is the most abundant glycoprotein in the virion, unlike the (E) protein which is present in minute amounts yet it is essential for coronavirus morphogenesis and envelope formation (Narayanan et al., 2000). Meanwhile, the (N) protein is present inside the virion complexed with the viral RNA to form the helical nucleocapsid structure (Risco et al., 1996). Another ORF (ORF1ab) encodes for a set of nonstructural proteins. The remaining ORFs (ORF3a, ORF6, ORF7a, ORF8, and ORF10) encode for a set of accessory proteins (Liu et al., 2014).

During the translation process from mRNA to protein, information is transmitted in the form of nucleotide triplets named codons. Amino acids are degenerate, having more than one codon representing each except for methionine (Met) and tryptophan (Trp). Thus, codons encoding the same amino acid are known as synonymous codons. Many studies on different organisms showed that synonymous codons are not used uniformly within and between different genomes. This phenomenon is called synonymous codon usage bias (SCUB) or codon usage bias (CUB) (Behura and Severson, 2012; Boël et al., 2016; Gu et al., 2004; Vicario et al., 2007). Hence, each organism has its preferred (optimal) codons, where a preferred codon is defined as a codon which is more frequently used in highly expressed genes than in less expressed genes (Wang et al., 2018). Two main forces shape the codon usage of an organism, mutation pressure and natural selection (Chen et al., 2014; Prabha et al., 2017; Zalucki et al., 2007). Other factors also known to influence the CUB are nucleotide composition (Palidwor et al., 2010), synonymous substitution rate (Marais et al., 2003), tRNA abundance (Rocha, 2004), codon hydropathy and DNA replication initiation sites (Huang et al., 2009), gene length (Duret and Mouchiroud, 1999), and expression level (Hiraoka et al., 2009). Since viruses rely on the tRNA pool of their hosts in the translation process, previous studies suggested that translational selection and/or directional mutational pressure act on the codon usage of the viral genome to optimize or deoptimize it towards the codon usage of their hosts (Burns et al., 2006; Cladel et al., 2008). Therefore, it is important to examine the composition of viral genes at the codon or nucleotide level to better understand the mechanisms driving virus-host relationships and virus evolution. In addition, it may assist in predicting the characteristics of newly emerging viruses (Sheikh et al., 2020). A study on Influenza A virus (IAV) (Goñi et al., 2012) suggested that understanding codon usage patterns in viruses may help in creating new vaccines using Synthetic Attenuated Virus Engineering (SAVE). By deoptimizing viral codons it is possible to attenuate a virus (Coleman et al., 2008). Another study reported that the replacement of natural codons with synonymous triplets with increased CpG frequencies gives rise to inactivation of poliovirus infectivity (Burns et al., 2009).

Due to the interest in CUB studies and their important contributions to fundamental research and various applications, we conducted a genome-wide codon usage analysis with the aim of characterizing the pattern of CUB in SARS-CoV-2 and defining the main force influencing it using 13 isolates from different geo-locations (countries). This may help in understanding the molecular evolution regarding the codon usage and nucleotide composition of SARS-CoV-2 and its adaptation to *Homo sapiens*. The information obtained in this study may also provide the basis for developing an attenuated SARS-CoV-2 vaccine.

## 2. Materials and Methods

### 2.1 Sequence data collection

Complete sequences of all the ten genes (N, E, S, M, ORF1ab, ORF3a, ORF6, ORF7a, ORF8, and ORF10) for 13 SARS-CoV-2 isolates were obtained from the NCBI virus portal (https://www.ncbi.nlm.nih.gov/labs/virus/vssi/#/) in FASTA format. Each of the 13 isolates was picked from a different country (USA, Pakistan, Spain, Vietnam, Italy, India, Brazil, China, Sweden, Nepal, Taiwan, South Korea, and Australia) according to the most recent date and data availability (at the time of this study). All information about the used isolates can be found in (Supplementary file 1). In this study, isolates were named by their geo-location (country).

### 2.2 Nucleotide composition analysis

The following compositional properties were calculated for the genes of the SARS-CoV-2 genomes: (i) the overall frequency of each nucleotide (A%, C%, U%, and G%); (ii) the frequency of each nucleotide at the third position of synonymous codons (A3%, C3%, U3%, and G3%); (iii) the GC and AU contents at the first, second, and third synonymous codon positions, as well as, the overall GC and AU contents; (iv) the G and C base incidences at the first (GC1), second (GC2) and third (GC3) synonymous codon positions; and (iv) the mean frequencies of nucleotides G and C at the first and second positions (GC12).

### 2.3 Synonymous Dinucleotide Usage (SDU)

A new index named Synonymous Dinucleotide Usage (SDU) has been developed (Lytras and Hughes, 2020) to implement a method to estimate the degree to which a host driven force acting on the dinucleotide level of viral genomes has skewed the synonymous codon usage of the protein sequence. To examine the occurrences of a given dinucleotide to the null hypothesis that there is an equal usage of synonymous codons. According to the SDU index, a coding sequence can have three distinct dinucleotide frame positions; frame 1 as the first and second nucleotide codon positions, frame 2 as the second and third nucleotide codon positions, and a bridge frame as the third nucleotide codon position and the first nucleotide from a downstream codon on the same coding sequence. The SDU values for frame 2 in all CDSs of SARS-CoV-2 isolates in this study were calculated using the following equation:

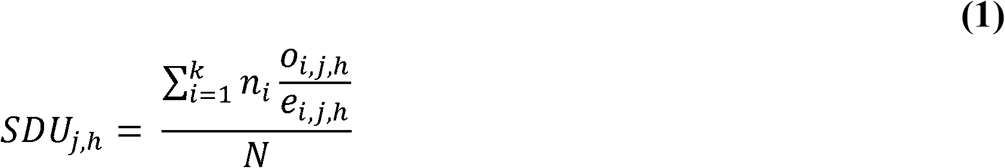

where *n_i_* is the number of occurrences of amino acid *i* in the sequence, *o_i,j,h_* is the synonymous proportion of dinucleotide *j* in frame position *h* for amino acid *i* observed in the sequence, *e_i,j,h_* is the synonymous proportion of dinucleotide *j* in frame position *h* for amino acid or amino acid pair *i* expected under equal synonymous codon usage, and *N* it the total number of amino acids in the sequence.

The result of the SDU directly reflects the overall synonymous dinucleotide representation in each frame of the tested CDSs. An SDU value of 1 indicates that, the representation of the dinucleotide in a given frame position is equal to the expected under the null hypothesis. An SDU value > 1 means that the dinucleotide is overrepresented in a given frame position compared to the excepted under the null hypothesis. Lastly, an SDU value between 0 and 1 shows an underrepresentation of the dinucleotide in the given frame position, compared to the representation assumed in the null hypothesis.

### 2.4 Relative Synonymous Codon Usage (RSCU)

Using the following equation (Sharp and Li, 1987):

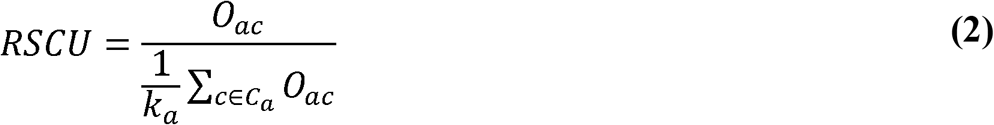

where *O_ac_* is the count of codon *c* for the amino acid *a*, and *k_a_* is the number of synonymous codons in amino acid *a* family, RSCU values were calculated.

An RSCU value of 1 indicates no codon usage bias as the observed frequency is equal to the expected frequency. RSCU values less than 1 indicate negative bias, and values of greater than 1 indicates positive bias. Accordingly, RSCU values are divided into 3 ranges; values ≤ 0.6 indicates underrepresented codons, values between 0.6 and 1.6 indicates randomly used codons, and values ≥ 1.6 indicates overrepresented or putative preferred codons (Duret and Mouchiroud, 1999; Liu et al., 2010; Mandal et al., 2020; Sharp and Li, 1987).

### 2.5 Effective number of codons (ENc)

The effective number of codons (ENc) determines the degree of preference for the unbalanced use of codons, regardless of the gene length and numbers of amino acids. ENc values range from 20, indicating extreme codon usage bias as only one synonymous codon is used for the corresponding amino acid, to 61, indicating no bias but the use of all possible synonymous codons equally. Therefore, the ENc value correlates negatively with the codon usage bias. Also, it is generally accepted that genes have significant codon usage bias when the ENc value is ≤ 35 (Comeron and Aguadé, 1998), (Wright, 1990). ENc was calculated using the following equation (Sun et al., 2013):

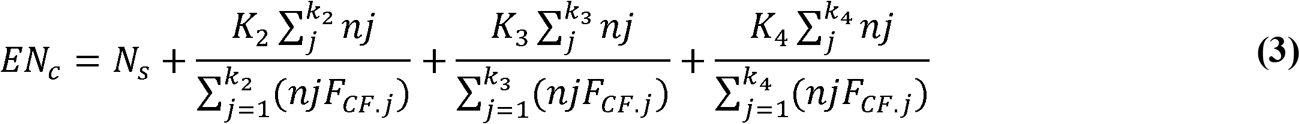

Where *N_S_* is the number of codon families with a single codon (e.g. Met and Trp). *K_i_* is the number of *i*-fold codon families. And, *F_CF.j_* is *F_CF_* for family *j* obtained from the following equation:

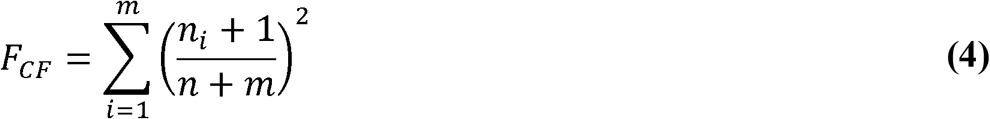

Where *n_i_* is the count of codon *i* in codon family of *m* synonymous codons.

An analysis of variance (ANOVA) was performed to determine whether the ENc values for each CDS of SARS-CoV-2 were significantly different or not.

### 2.6 ENc-GC3 plot

To determine whether the codon usage of given genes is solely due to mutational pressure or selectional pressure, an ENc-plot was drawn where the expected ENc values from GC3s (denoted by `S’) were determined according to the following equation (Novembre, 2000; Wright, 1990):

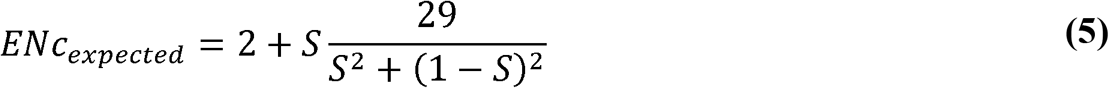

Using the above equation, the expected fitting curve of ENc values was drawn then ENc versus GC3s values for each coding region are plotted. If the distribution of the plotted genes is along/near the curve, then the codon usage bias is assumed to be affected only by mutation. If the distribution of the plotted genes is below the curve, then the codon usage bias is assumed to be affected by selection and other factors (Liu et al., 2012).

### 2.7 Neutrality Plot

Neutrality plot is used to estimate the effect of mutation pressure and natural selection on codon usage bias. In this study, the GC contents at the first, second and third codon positions (GC1, GC2, GC3, respectively) of the 13 SARS-CoV-2 isolates were analyzed. Then, GC12 representing the average GC content at the first and second codon positions of each isolate was obtained. Both GC12 and GC3 values were used for neutrality plot analysis. If the correlation between GC12 and GC3 is statistically significant and the slope of the regression line is close to 1, mutation pressure is assumed to be the main force structuring codon usage bias. Conversely, natural selection would have higher odds leading to a narrow distribution of GC content and a lack of correlation between GC12 and GC3 (Jia and Xue, 2009; Song et al., 2017; Wu et al., 2015).

### 2.8 Grand Average of Hydropathicity (GRAVY) and Aromaticity (AROMO)

The GRAVY value is calculated by dividing the sum of hydropathy values of all amino acids in a sequence by the number of residues (Kyte and Doolittle, 1982). GRAVY values range between −2.0 and +2.0, with positive values indicating protein hydrophobicity, while negative values indicate protein hydrophilicity.

The AROMO value reflects the recurrence of aromatic amino acids such as tryptophan, phenylalanine, and histidine in a given amino acid sequence.

### 2.9 PR2-bias Plot Analysis

The Parity Rule 2 (PR2) bias was calculated based on the values of A/U-bias ‘A3 / (A3 + U3)’ and G/C-bias ‘G3 / (G3 + C3)’, they were plotted as the ordinate and abscissa, respectively. The centre of the plot is the place where both coordinates are 0.5, as well as, A = U and G = C according to Chargaff’s rule (PR2) (Rapoport and Trifonov, 2013).

### 2.10 Codon Adaptation Index (CAI)

Codon adaptation index (CAI) uses a reference set of highly expressed genes (e.g. ribosomal genes), this measure is an indicator of gene expression levels and natural selection; it ranges from 0 to 1 with higher values indicating stronger bias with respect to the reference set, therefore this method is an indicator of selection for a bias toward translational efficiency (Ran and Higgs, 2012; Sharp and Li, 1987). Codon adaptation index (CAI) was calculated by the equation given by (Lee, 2018; Ran and Higgs, 2012):

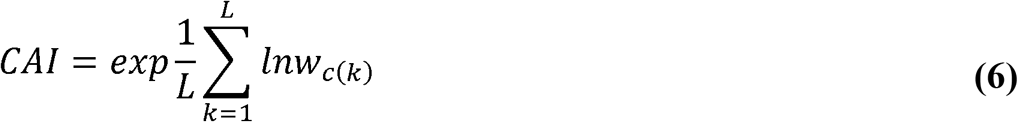

where *L* is the count of codons in the gene and *w_c(k)_* is the relative adaptiveness value for the *k-th* codon in the gene.

Twelve genes with the highest level of expression in the lung tissues for human were collected from the human protein atlas project database (https://www.proteinatlas.org/humanproteome/tissue/lung). Then, CAI values for the 13 isolates were calculated based on those 12 genes as a reference set.

### 2.11 tRNA Adaptation Index (tAI)

The tRNA Adaptation Index (tAI) is a measure of tRNA usage by a CDS considering the fact that the tRNA gene copy number of some genomes has a high positive correlation with tRNA abundance within a cell, as well as, codon usage preferences (dos Reis et al., 2004). Recently, a novel approach was developed that estimates species-specific tAI wobble weights by optimizing the correlation between a measure of codon usage bias and tAI, without the need to utilize the gene expression measurements of the test organism (based on the CUB) (Renana and Tamir, 2014). The frequencies of tRNAs of *Homo sapiens* was obtained from the GtRNAdb (http://gtrnadb.ucsc.edu/) (Chan and Lowe, 2009).

### 2.12 Relative Codon Deoptimization Index (RCDI)

The relative codon deoptimization index (RCDI) is a comparative measure that is used to determine the codon deoptimization trends by comparing the similarity in codon usage between a given coding sequence and a reference genome (Mueller et al., 2006). If the tested coding sequence of a virus has a similar codon usage pattern to the human genome, the RCDI value would be closer to or equal 1, which may indicate a high translation rate, as well as, a better adaptation to the host. RCDI values greater than 1 indicates a deviation from the normal human codon usage bias (deoptimized viral genome). In this study, the synonymous codon usage patterns for *Homo sapiens* were used as a reference and obtained from the CUD. Non-synonymous codons and termination codons were excluded from the analysis. The RCDI was calculated using the following equation (Mueller et al., 2006):

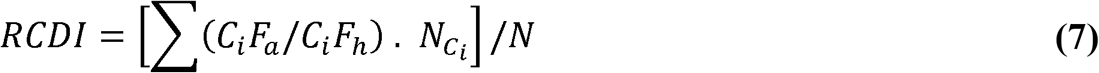

Where *C_i_F_a_* is the observed relative frequency of codon *i* to other synonymous codons for the same amino acid in the tested CDS; *C*_i_*F_h_* is the normal relative frequency of codon *i* for a specific amino acid in the reference genome sequence; *N_Ci_* is the number of incidences of codon *i* in the CDS; and *N* is the total number of codons in the CDS.

### 2.13 Phylogenetic Analysis

A dendrogram was constructed using 13 complete genomic sequences of 13 SARS-CoV-2 isolates obtained from different geo-locations (Countries). Evolutionary relationships were inferred by using maximum-likelihood statistical method with general time-reversible (GTR) model implemented in MEGA-X software (v10.1) (Tamura et al., 2011). The bootstrap method with 1000 replicates was used to test the reliability of the phylogenetic tree. For each isolate, the following data are given: Species and country of origin.

### 2.14 Software and Statistical analysis

Spearman’s rank correlation and linear regression analyses were performed using R Language (R Core Team, 2016). Different R packages as vhcub, SeqinR, ggplot2 and stats (Anwar et al., 2020; Charif and Lobry, 2007; R Core Team, 2018; Wickham, 2016) were used to calculate various CUB indices and to draw the graphs in this study. As well as a python package named CAI (Lee, 2018) was used to estimate the CAI for the tested viral isolates. The RCDI was calculated using a python script (available upon request). tAI was calculated using stAIcalc (Sabi et al., 2017). The cluster analysis (Heat map) was performed using CIMminer (https://discover.nci.nih.gov/cimminer/) based on the RSCU values obtained from the tested isolates. Multiple sequence alignments for the whole genome of the 13 SARS-CoV-2 was done with MAFFT software (v7.450) and the phylogenetic analysis was performed with MEGA-X software (v10.1).

## 3. Results

### 3.1 Nucleotide and Dinucleotide Composition Analysis

The analysis of nucleotide content of the coding sequences (CDSs) in all tested SARS-CoV-2 isolates showed A and U nucleotides richness, in comparison to G and C nucleotides, as well as, AU bias in all codon positions (Supplementary file 2, and Figure 1). The average usage (%) of U was the highest (33.24 ± 4.75%) among the four nucleotides; A (28.91 ± 3.74%) showed the highest usage after U, followed by C (19.50 ± 2.88%), while G showed the lowest usage (18.32 ± 2.22%). A similar trend of nucleotides content at the third positions of synonymous codons was observed (U3% > A3% > C3% > G3%); where U3% (40.07 ± 4.51%) and A3% (27.68 ± 3.83%) were higher than C3% (18.37 ± 3.92%) and G3% (13.86 ± 2.47%). The mean AU and GC values were 62.16 ± 4.78% and 37.83 ± 4.78%, and the mean AU3% and GC3% values were 67.79 ± 5.01% and 32.20 ± 5.014%, respectively.

**Figure 1.**
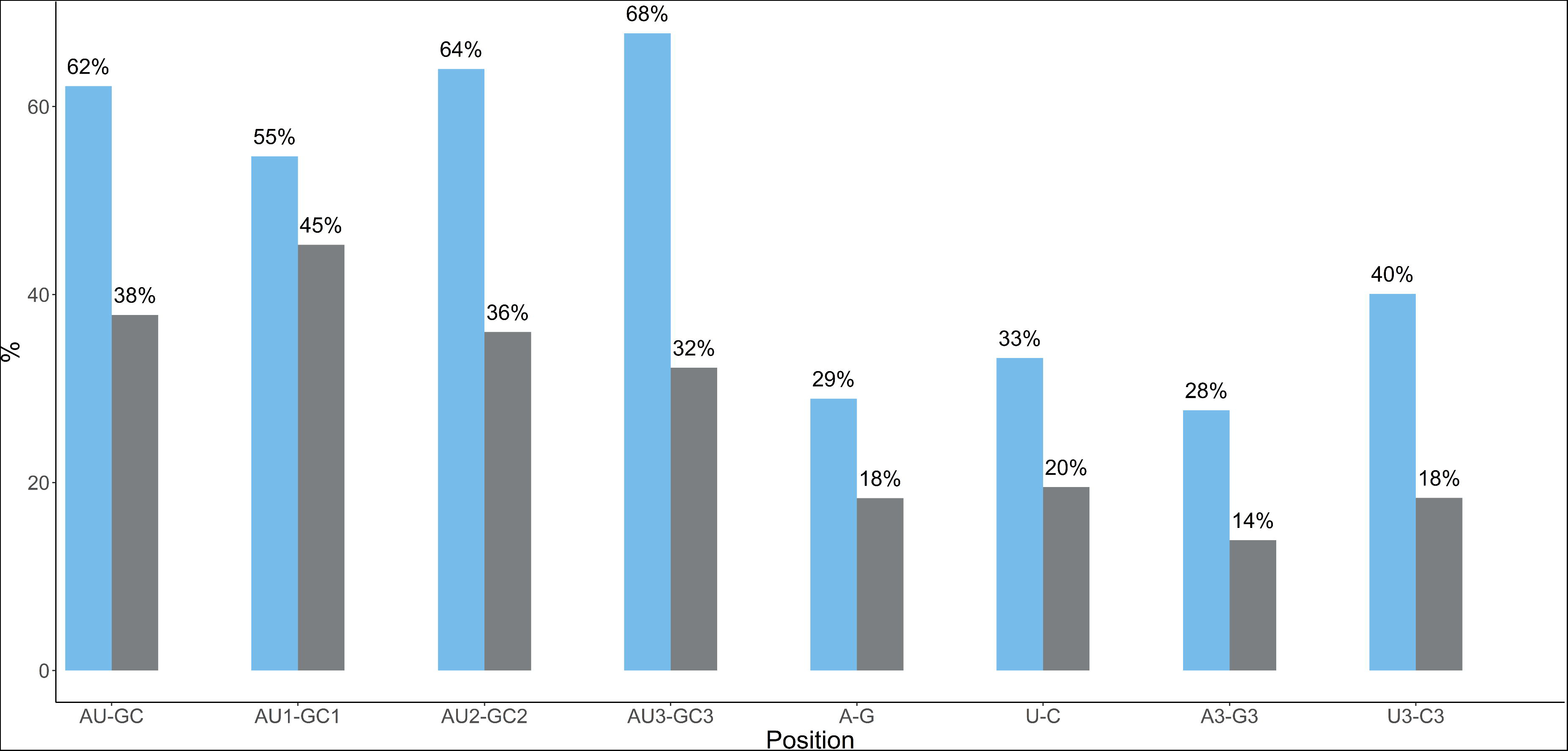
The nucleotide composition of SARS-CoV-2. From left to right it includes the following: (i). overall AU and GC contents. (ii). AU and GC contents and the first, second, and third codon positions. (iii). over all frequency of each nucleotide (A%, U%, G%, C%). (iv). the frequency of each nucleotide at the third codon position (A3%, U3%, G3%, C3%)

Synonymous Dinucleotide Usage (SDU) was employed to reveal underrepresented and overrepresented dinucleotides in the second and third codon positions (frame 2). The average SDU values for each dinucleotide in all examined isolates were reported by a bar plot (Figure 2). Seven dinucleotides exhibited overrepresentation (SDU > 1), in descending order (CpU, UpU, ApA, GpU, CpA, UpA, and ApU). All these dinucleotides are A/U-ended (A-ended: 3, U-ended: 4) reflecting the A/U-bias of SARS-CoV-2 genomes at the third codon positions. On the other hand, nine dinucleotides showed underrepresentation (SDU < 1), in ascending order (CpC, GpG, ApG, CpG, UpG, GpC, UpC, ApC, and GpA). Eight out of the nine underrepresented dinucleotides were G/C-ended (G-ended: 4, C-ended: 4).

**Figure 2.**
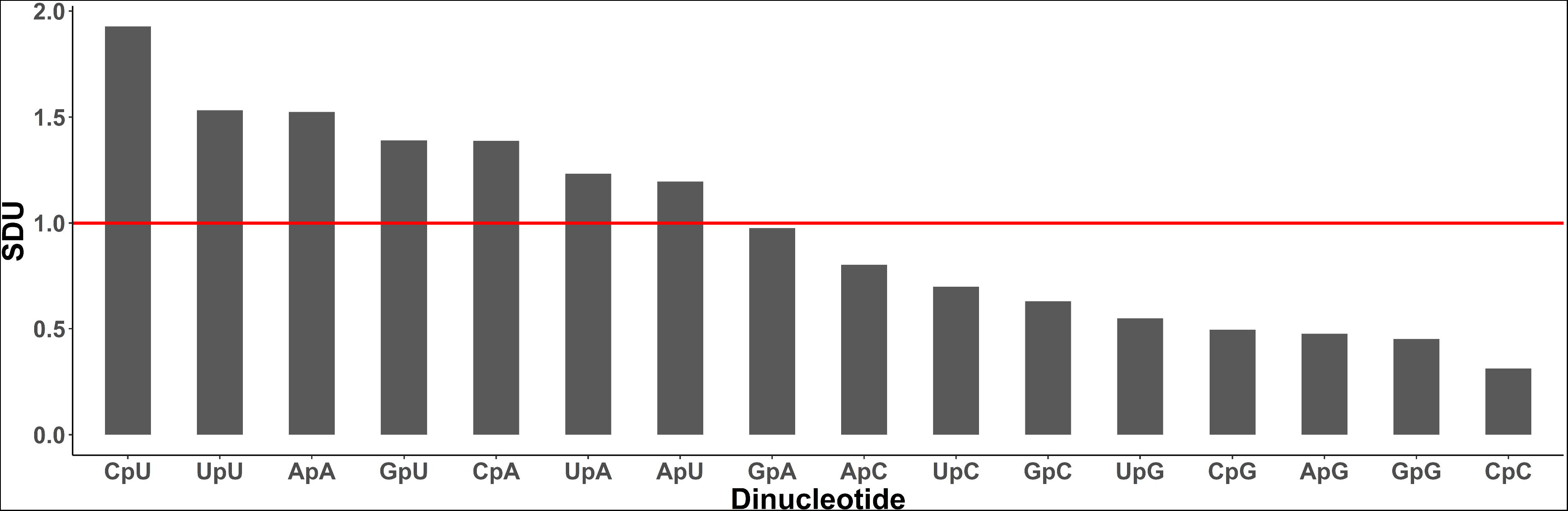
The average synonymous dinucleotide usage (SDU) values for each dinucleotide from all examined isolates are estimated and plotted. The x-axis represents each dinucleotide and the y-axis shows the average SDU for each dinucleotide. The red line signifies the cut off (SDU > 1) of dinucleotide to be overrepresented.

### 3.2 Relative Synonymous Codon Usage (RSCU) and Cluster (Heat map) analyses

In order to determine to what extent A/U-ended codons are preferred, and the patterns of synonymous codon usage, RSCU values were calculated for the complete coding region of all 13 SARS-CoV-2 isolates. Among the 18 amino acids used in the analysis, eight common overrepresented codons were found in all tested isolates for the following amino acids (Arg, Val, Ser, Ala, Pro, Thr, Leu, and Gly) with their corresponding average RSCU values (2.47, 2.03, 2.00, 1.82, 1.72, 1.70, 1.68, 1.65, respectively) (Figure 3, and Supplementary file 3). The amino acid Arginine (Arg) over biased with AGA codon, Valine (Val) over biased with GUU, Serine over biased with UCU, Alanine (Ala) over biased with GCU, Proline (Pro) over biased with CCU, Threonine (Thr) over biased with ACU, Leucine (Leu) over biased with CUU, and Glycine (Gly) over biased with GGU. All previously mentioned overrepresented codons are A/U-ended (U-ended: 7, A-ended: 1), and four of them had the CpU dinucleotide in the second and third codon positions which is the most represented dinucleotide in frame 2 according to the dinucleotide composition analysis. On the other hand, almost all underrepresented codons (RSCU <1) are G/C-ended.

**Figure 3.**
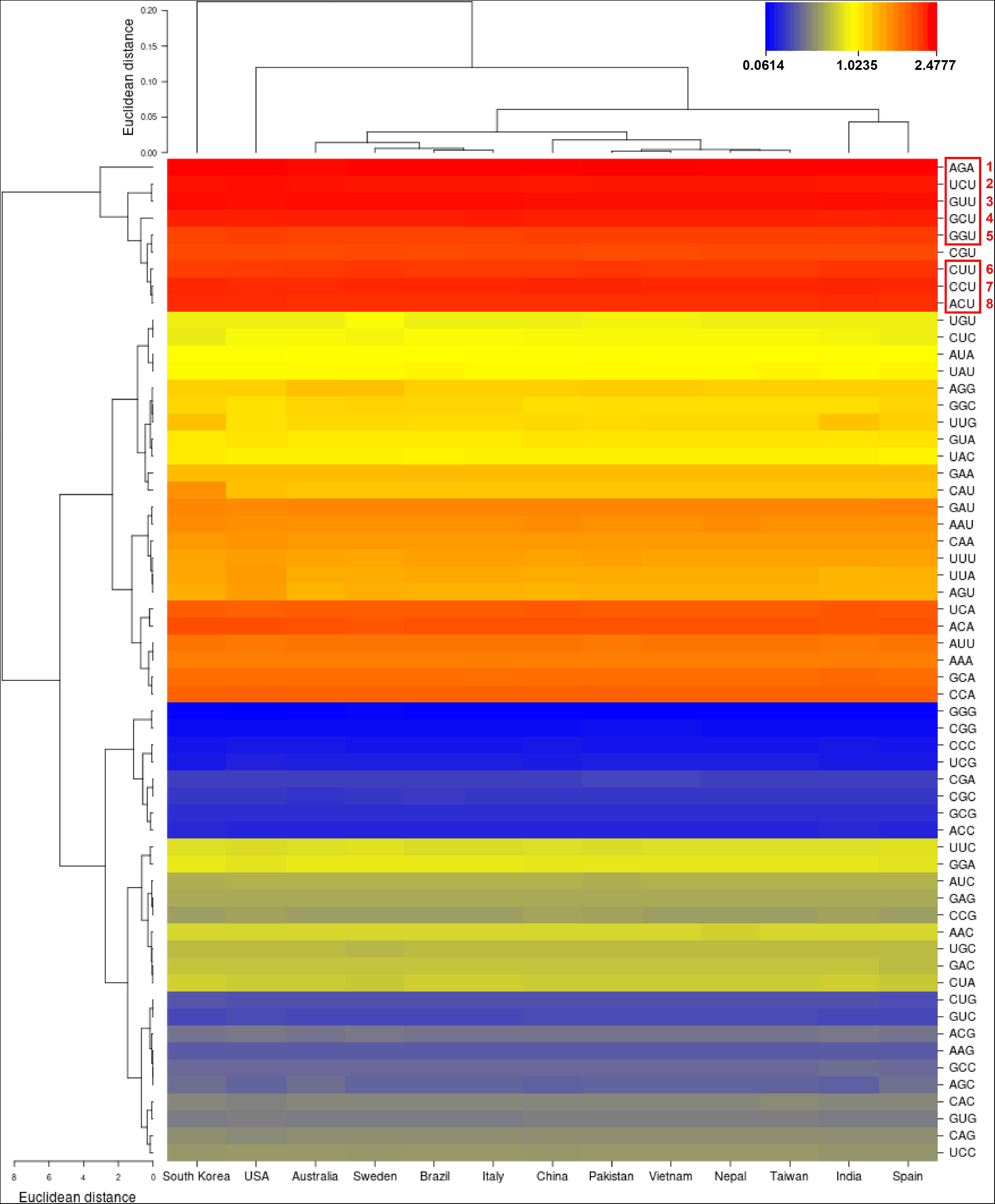
Cluster (Heat map) analysis based on the RSCU values of the 13 SARS-CoV-2 isolates complete coding regions. Higher RSCU values representing more frequent codon usage depicted in red colour, and lower RSCU values depicted in blue colour

To illustrate the similarities of synonymous codon usage patterns across all 13 tested isolates, a cluster (heat map) analysis was performed based on the average RSCU values of their CDSs (Figure 3). All isolates had similar synonymous codon usage patterns with slight differences in their randomly used or biased codons (RSCU values from 0.6 to 1.6). According to the cluster analysis, the following isolates showed similar synonymous codon usage patterns: (i) India and Spain, (ii) China, Pakistan, Vietnam, Nepal, and Taiwan, (iii) Australia, Sweden, Brazil, and Italy. Meanwhile, both, South Korea and USA isolates grouped themselves from the remaining ones reflecting their unique synonymous codon usage patterns. Interestingly, cluster analysis results were comparable with the phylogenetic analysis suggesting that patterns of synonymous codon usage were able to reflect the evolutionary relationships between tested isolates (Figure S1).

### 3.3 Effective number of codons (ENc) and ENc-GC3 plot

In theory, ENc correlates negatively with CUB. For the complete coding region of all tested isolates, the average ENc value was 50.38 ± 0.0042 (Supplementary file 4). The ENc values of individual genes was different, ranging from 45.78 to 54.09 with an average of 50.38 ± 2.38 (Supplementary file 2). The ORF10 gene had the highest ENc value in all isolates 54.09 ± 0.00, while the ORF1ab gene had the lowest ENc value 45.78 ± 0.02. The analysis of variance (ANOVA) showed that the average ENc values of the individual genes of SARS-CoV-2 were significantly different (*P* <.001). Tukey HSD test was performed at a 95% confidence level and revealed that there was a statistically significant difference in the average ENc values between all SARS-CoV-2 genes used in the analysis (N, E, S, M, ORF1ab, ORF3a, ORF6, ORF7a, ORF8, and ORF10). Overall, ENc results suggest that the codon usage of SARS-CoV-2 is slightly biased (ENc > 35).

To better understand the relations between SARS-CoV-2 genes composition and codon usage bias, an ENc-GC3 plot was drawn. If the codon usage pattern is only affected by GC3 resulting from mutation pressure, the data points should be just on or near the expected curve drawn by equation (5) and shown in (Figure 4). In the figure, though some points fell onto the expected curve, the majority were far from the curve. The results indicated that codon usage bias tended to be stronger when GC contents were lower, and vice versa.

**Figure 4.**
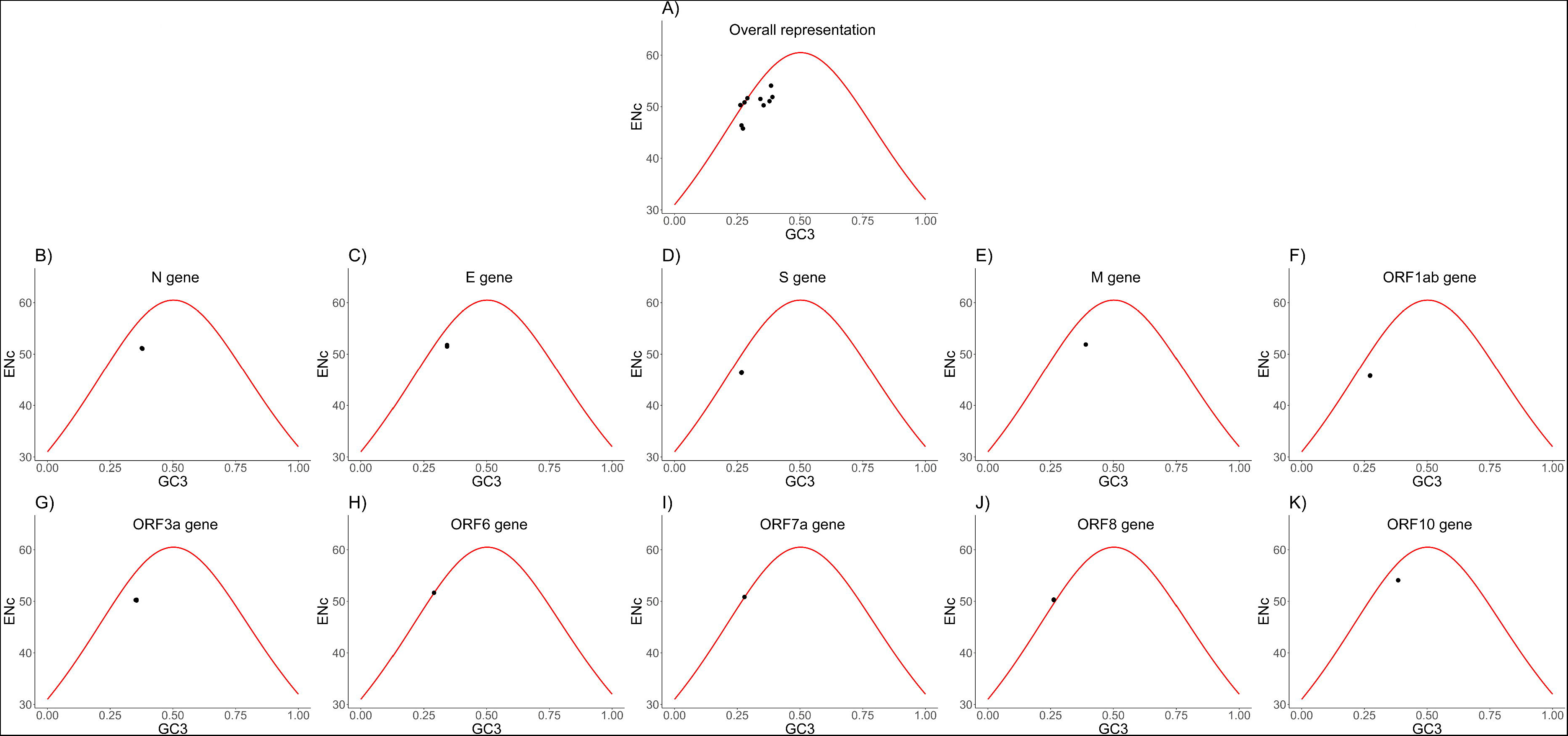
ENc-GC3 plots of different genes of SARS-CoV-2. The solid line represents the expected curve when codon usage bias is only affected by mutation pressure. **(A)** ENc-GC3 plot for all genes; **(B—K)** for individual genes (N, E, S, M, ORF1ab, ORF3a, ORF6, ORF7a, ORF8, and ORF10, respectively).

### 3.4 Parity rule 2 (PR2) bias analysis

PR2 bias plots are informative when A/U-bias and G/C-bias at the third codon position in four-codon amino acids of individual genes are plotted. If codon usage bias is only caused by mutation pressure, AU or GC should be used proportionally among the degenerate codon groups in a gene. Contrarily, natural selection for codon choice would not necessarily cause proportional use of G and C (A and U) (Sueoka and Kawanishi, 2000). The four-codon amino acids are alanine, arginine (CGA, CGU, CGG, CGC), glycine, leucine (CUA, CUU, CUG, CUC), proline, serine (UCA, UCU, UCG, UCC), threonine, and valine. The average A/U-bias and G/C-bias values were 0.38 ± 0.061 and 0.39 ± 0.222, respectively (Figure 5). Therefore, U will be preferred over A and likewise C will be preferred over G, in other words, pyrimidines are preferred over purines in SARS-CoV-2.

**Figure 5.**
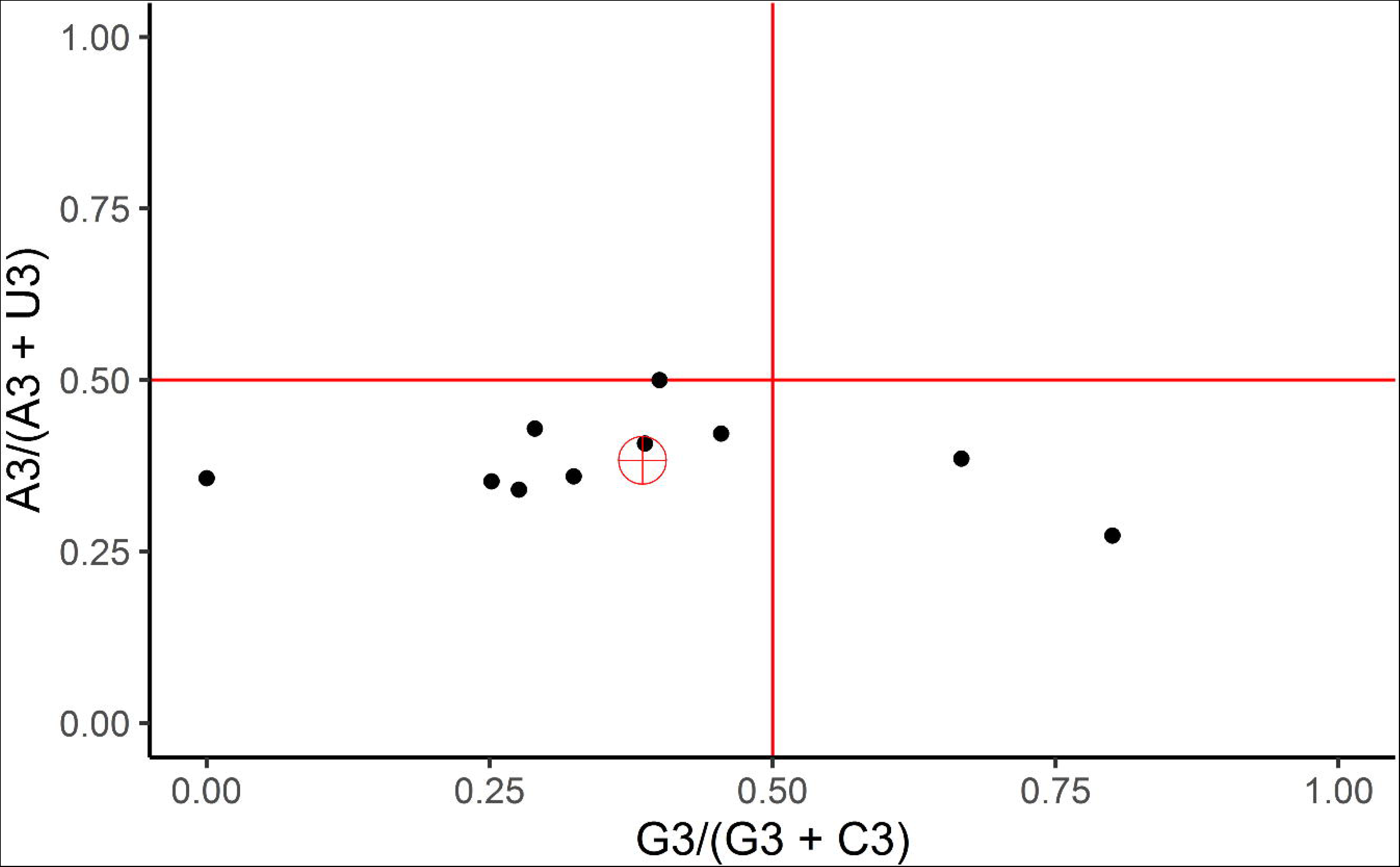
PR2-bias plot [A3/(A3◻+◻U3) against G3/(G3◻+◻C3)] of SARS-CoV-2 four-fold degenerate codons.

### 3.5 Neutrality analysis

ENc-GC3 plot and parity analyses reflected the main factors influencing the codon usage bias of SARS-CoV-2, but they did not estimate precisely which of mutation pressure or natural selection was more important. Then a neutrality plot was constructed based on the average GC3 and GC12 contents of SARS-CoV-2 genes (Figure 6). The plot revealed a narrow distribution range (0.262-0.39) of GC3 contents, while GC12 contents were relatively wide (0.274-0.518). The correlation between GC1 and GC2 was strong (r = 0.76, *P* =.016), meanwhile, neither GC1 nor GC2 showed significant correlation with GC3 (r = 0.14, *P* =.69), reflecting that codon usage is affected by mutation pressure to a very limited degree. In addition, the slope of the regression line revealed that mutation pressure only accounted for 18.4% of SARS-CoV-2 codon usage, while natural selection together with some other factors accounted for the remaining 81.6%.

**Figure 6.**
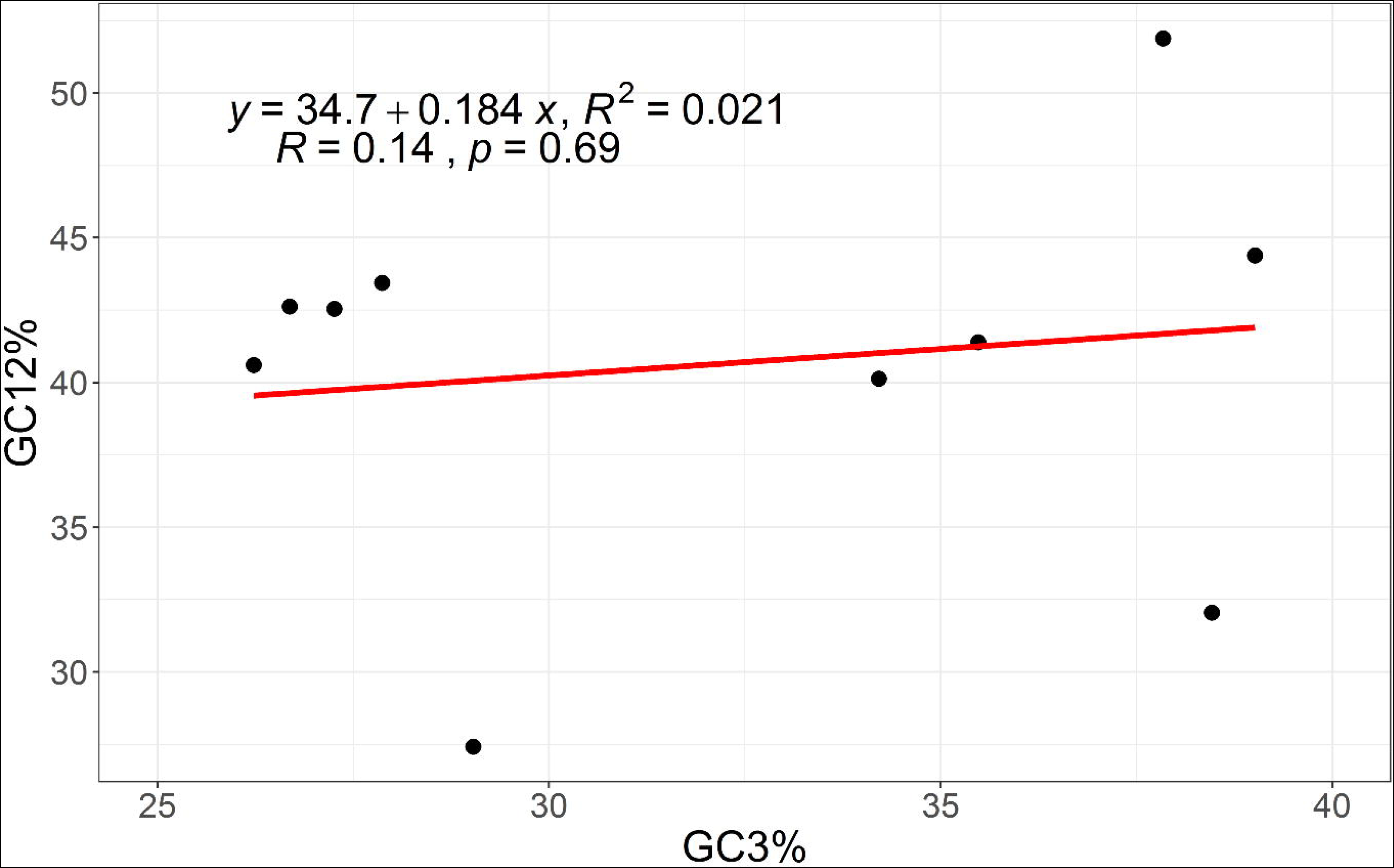
Neutrality plot analysis of GC12 and GC3 contents. GC12 frequencies were plotted against GC3 frequencies. The y-axis (GC12) refers to the average GC frequency at the first and second codon positions. The x-axis (GC3) refers to the GC frequency at the third codon position. The slope value indicates the mutational pressure.

### 3.6 Codon Adaptation Index (CAI)

To determine the degree of adaptation of SARS-CoV-2 codon usage to *Homo sapiens*, the CAI values for the complete coding region and individual genes of each isolate was calculated using the most expressed genes in human lung tissues (the reservoir of viral host cells) as a reference set. The average CAI value for the complete coding region was 0.54 ± 0.0003 (Supplementary file 4). In addition, the CAI values of different genes ranged from 0.48 to 0.62 (Figure 7, and Supplementary file 2). The N, ORF7a, and S genes had the highest average CAI values (0.62 ± 0.0008, 0.58 ± 0.0000, 0.56 ± 0.0002, respectively) among other genes, while the E gene had the lowest CAI value (0.48 ± 0.001).

**Figure 7.**
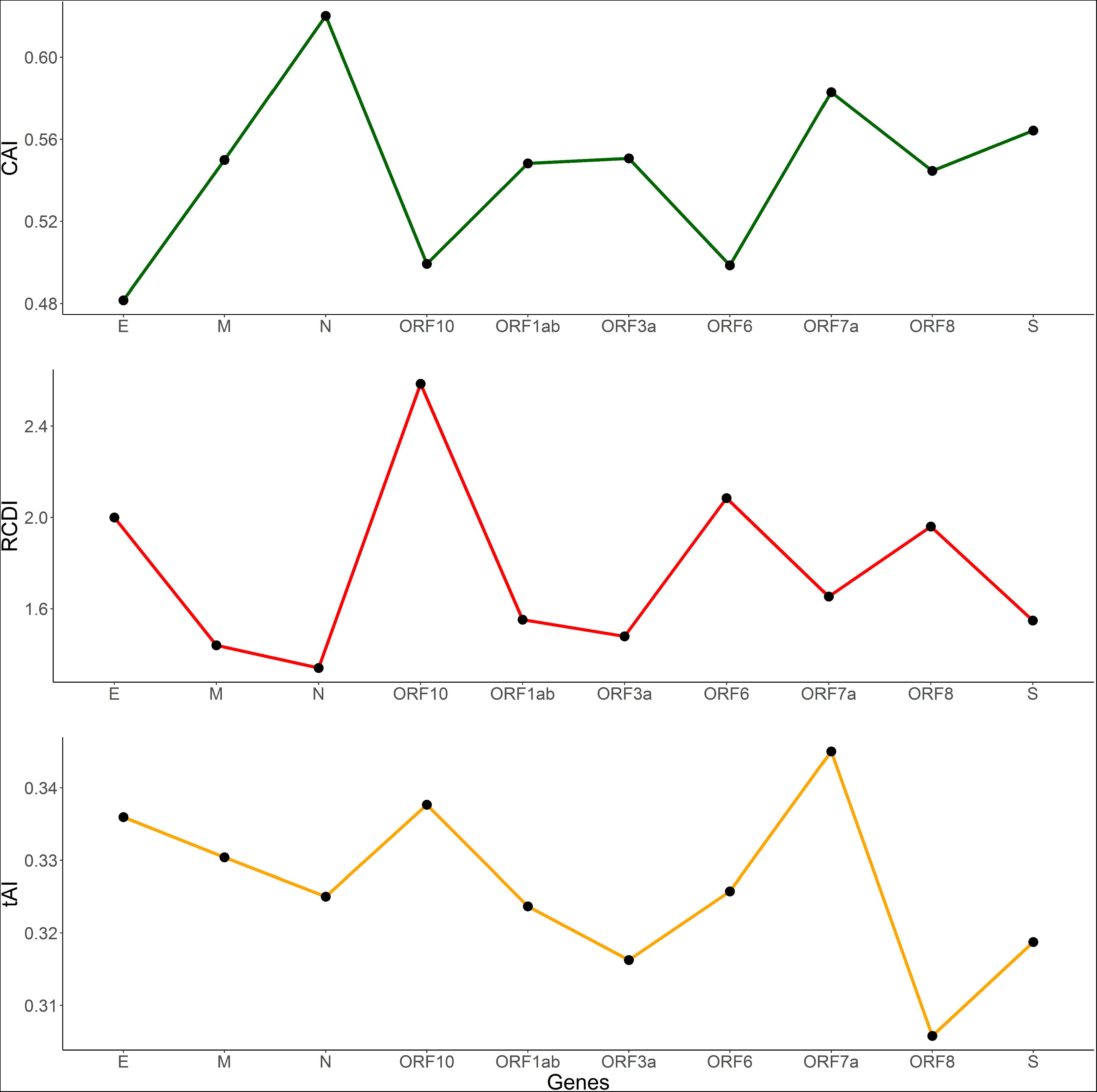
The Codon adaptation index (CAI), relative codon deoptimization index (RCDI), and tRNA adaptation index (tAI) analyses of SARS-CoV-2 genes in relation to *Homo sapiens*.

### 3.7 Relative Codon Deoptimization Index (RCDI)

The relative codon deoptimization index (RCDI) was measured by comparing the codon usage of SARS-CoV-2 to *Homo sapiens*. The lower the RCDI value, the higher the virus adaptation to the host. The average RCDI value for the complete coding region across all isolates was 1.76 ± 0.003 (Supplementary file 4). Within genes, the RCDI values ranged from 1.34 to 2.58 with an average of 1.76 ± 0.37. The lowest RCDI value was 1.34 ± 0.0004 for the N gene, whereas the highest value was 2.58 ± 0.00 for the ORF10 gene (Figure 7, and Supplementary file 2). The Tukey HSD test at 95 confidence level showed statistically significant differences in the RCDI values between all genes except ORF1ab and S.

### 3.8 tRNA Adaptation Index (tAI)

In order to assess the influence of translational selection on SARS-CoV-2 codon usage bias, the tRNA adaptation index (tAI) was calculated for the complete coding region and individual genes of tested isolates. The average tAI value for the complete coding region across all isolates was 0.33 ± 0.016 (Supplementary file 4). In addition, different genes showed different tAI values ranging from 0.31 ± 0.016 (ORF8) to 0.35 ± 0.014 (ORF7a) with an average of 0.33 ± 0.011 (Figure 7, and Supplementary file 2), indicating distinct intragenic translation efficiency patterns.

## 4. Discussion

### 4.1 A/U Richness of the SARS-CoV-2 Genome

The nucleotide compositions in the genes of SARS-CoV-2 were analyzed and correlated with the RSCU patterns. The SARS-CoV-2 genome is found to be rich in AU as compared to GC overall content, as well as, in all codon positions. In addition, all overrepresented codons were A/U-ended, whereas overwhelming majorities of underrepresented codons were G/C-ended.

### 4.2 Compositional Constraints Versus Natural Selection

During the course of evolution, many vertebrate RNA viruses tended to exhibit reduced frequencies of CpG-containing codons, such observation can be explained by a host specific selection pressure acting against CpG dinucleotides driven by their innate immune response (Belalov and Lukashev, 2013). In this study, CpG dinucleotide was among the most four underrepresented dinucleotides in frame 2 (second and third codon positions) of SARS-CoV-2 coding regions. Moreover, seven out of all eight codons containing CpG dinucleotides (CGG, UCG, GCG, CGC, CGA, ACG, CCG) were underrepresented (RSCU < 0.65). Therefore, CpG depletion caused by selection pressure contributed significantly in shaping the codon usage patterns in SARS-CoV-2.

UpA dinucleotide underrepresentation is consistent in most living organisms including viruses (Shackelton et al., 2006). Higher frequencies of UpA nucleotides are often correlated with mRNA instability and susceptibility to degradation by cytoplasmic RNAses (Duan and Antezana, 2003). In addition, two out of the three stop codons (UAA and UAG) contain UpA dinucleotide, thus UpA deprivation may help in avoiding non-sense mutations. Surprisingly, in SARS-CoV-2, UpA was not underrepresented, and the reason might be attributed to the A/U richness of SARS-CoV-2 genome. Furthermore, all six UpA-containing codons (AUA, CUA, UUA, GUA, UAC, UAU) were not underrepresented with RSCU values ranging from 0.7 to 1.2, indicating the unbiased use of them. Consequently, this observation highlighted the role of compositional constraints in shaping the codon usage patterns of SARS-CoV-2, which dominated over the effect of the selective force that causes UpA depletion.

### 4.3 The disparity in CUB Between SARS-CoV-2 and *Homo sapiens*

The patterns of CUB of SARS-CoV-2 showed almost absolute antagonism to that of *Homo sapiens*, as both of them were biased to A/U- and G/C-ending codons, respectively (Supplementary file 5). Such phenomenon was reported in many other viruses such as henipavirus, poliovirus, and hepatitis A virus (Kumar et al., 2018; Mueller et al., 2006; Sanchez et al., 2003). Codon usage is known to influence translation efficiency (and/or translation accuracy) and protein folding, as the rapid production of proteins due to higher translation rate may lead to incorrect folding and aggregation of viral protein products. Therefore, this disparity in codon usage between viruses and their hosts may exist to enable proper folding of viral protein products (Hu et al., 2011). Previous studies focused on the fact that for viruses to be adapted to their hosts, they should evolve into a direction where both CUB are similar to facilitate viral expression, and any deviation is attributed to genetic drift and mutational bias (Jenkins and Holmes, 2003). However, a recent study highlighted the effect of previously unrecognized complexity in the coevolution of virus-host pairs where over similarity of CUB between viruses and their hosts is detrimental, because it will impede host translation (Chen et al., 2020). Hence, a convenient deviation between the CUB of viruses and their hosts is of particular importance to host fitness, and so successful life cycle of the virus. Accordingly, natural selection on viral expression is unlikely to be unidirectional to the end where viral and host CUB are overly similar, because many regulatory elements such as CUB and promoter sequences are involved in determining viral expression (Chen et al., 2020), further supporting the role of natural selection in shaping the codon usage of SARS-CoV-2.

### 4.4 SARS-CoV-2 Genome is Slightly Biased

The ENc values were calculated in the complete coding region of SARS-CoV-2 and also in individual genes to estimate the overall codon usage bias. The overall codon usage for the complete coding region is found to be slightly biased across all tested isolates with an average of 50.38 ± 0.0042. A very similar pattern of slight CUB has been reported in a previous study on 50 different RNA viruses, as they had ENc values ranging from 38.9 to 58.3 with an average of 50.9 (Jenkins and Holmes, 2003). Furthermore, many other studies on different RNA viruses for different hosts showed ENc values in the same range, as in H1N1pdm IAV (ENc = 52.5) (Goñi et al., 2012), Equine Infectious Anemia Virus (ENc = 43.61) (Yin et al., 2013), Bovine Viral Diarrhea Virus (ENc = 50.91) (Wang et al., 2010), and Classical Swine Fever Virus (ENc = 51.7) (Tao et al., 2008). On the other hand, different SARS-CoV-2 genes exhibited significantly different bias (*P* >.001), which can be explained by various selection pressures acting over their corresponding proteins. Overall, the tendency of RNA viruses to maintain such a slightly biased codon usage might have facilitated their ability to cross species barrier. Also, it might have a selective advantage for their efficient replication by avoiding the competition with their hosts for the synthesis machinery (Kumar et al., 2018), which is consistent with the observation that SARS-CoV-2 and *Homo sapiens* have significantly different CUB (in terms of most preferred codon for each amino acid).

### 4.5 SARS-CoV-2 Codon Usage is Shaped by Multiple Factors

Codon usage bias (CUB) is affected by many factors, and two generally accepted major forces are mutation pressure and natural selection. In order to define the main force and other factors involved in forming the CUB of SARS-CoV-2 complete coding region, we followed a stepwise manner and performed ENc-GC3, PR2 bias, and neutrality plot analyses. In ENc-GC3 plot when codon usage pattern is only affected by GC3 composition indicating mutation pressure, all ENc values should be just on the expected curve. The ENc values of all ten SARS-CoV-2 genes used in the analysis were below the expected curve except for three genes (ORF6, ORF7a, and ORF8), indicating that mutation pressure may be a factor in the formation of SARS-CoV-2 codon usage bias, but other independent factors such as natural selection strongly affected the codon usage bias pattern reflecting their importance over the mutation pressure. Then, a PR2 bias plot was generated to confirm that the pattern of codon usage is not only affected by mutation pressure. The results showed a clear bias in the third codon position of four-codon amino acids in SARS-CoV-2 genes. This observation also indicated that natural selection pressure might play a major role in SARS-CoV-2 codon usage bias, and mutation pressure would be a minor factor. In order to accurately estimate the effect of mutation pressure and natural selection on the CUB, a neutrality plot was drawn. When the CUB is only affected by mutation pressure, the quantitative relation between GC3 and GC12 should be nearly equal, and the slope would be 1. In this study, GC1 and GC2 showed no significant correlation with GC3 and the slope of regression line revealed that mutation pressure only accounted for 18.4% of SARS-CoV-2 codon usage, while natural selection together with some other factors accounted for the remaining 81.6%. Accordingly, natural selection played a very important or even a dominant role in the formation of SARS-CoV-2 codon usage. Furthermore, to determine whether codon choices were influenced by hydropathicity, a correlation analysis was performed between ENc, GRAVY, and AROMO (Table 1). The results showed no correlation between codon usage and aromaticity; however, a significant positive correlation was found between codon usage and hydrophobicity of SARS-CoV-2 coding sequences reflecting its effect on CUB. The link between hydropathy and codon usage may be caused by the fact that the expressed sequences are hydrophilic just because they accomplish their function in the aqueous media of the cell (Zhong et al., 2007). Furthermore, a correlation analysis was performed between the average tAI and ENc values of SARS-CoV-2 genes. The analysis revealed a positive significant correlation (r = 0.66, P =.04) (Table 1), which implies that translational selection significantly influenced the pattern of codon usage in SARS-CoV-2. These results combined confirm the dominance of natural selection over mutation pressure in shaping the CUB of SARS-CoV-2.

**Table 1.**
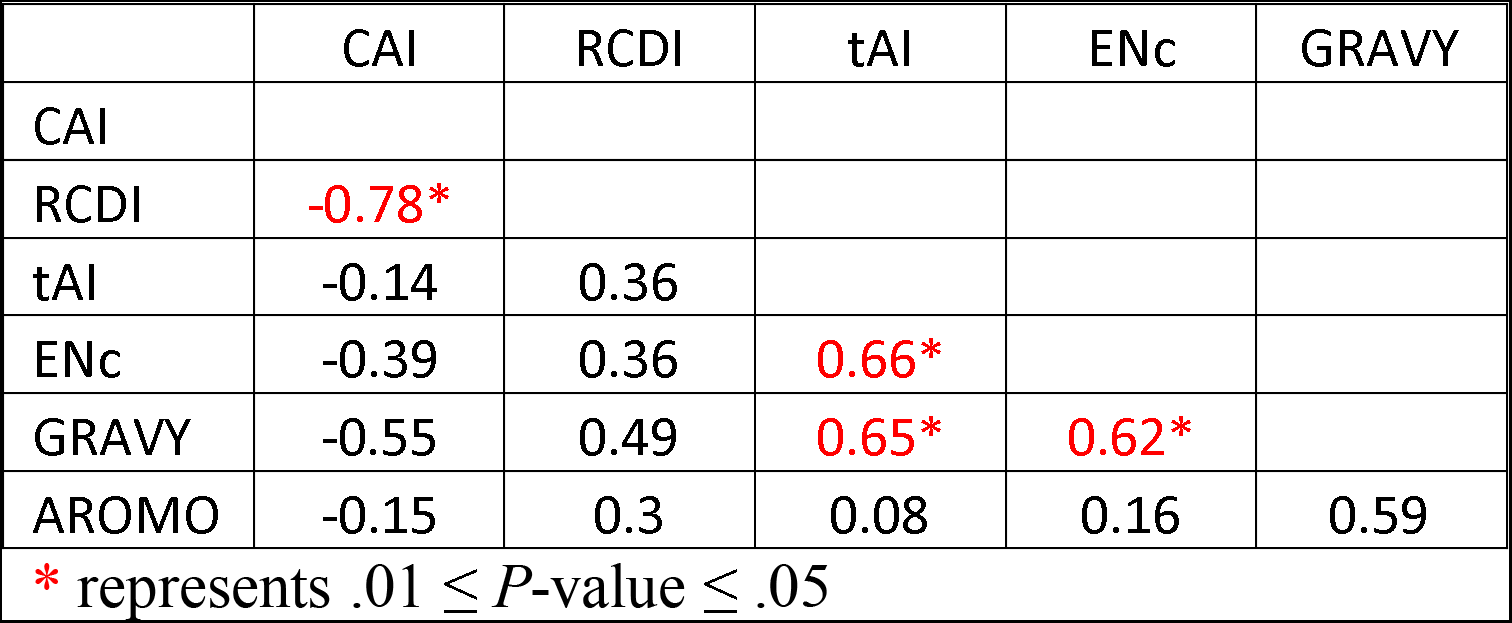
Correlation analysis between ENc, CAI, GRAVY and AROMO values.

### 4.6 Moderate Adaptation of SARS-CoV-2 to *Homo sapiens*

In order to determine to what extent SARS-CoV-2 is adapted to *Homo sapiens*, two adaptation indices were used (CAI and RCDI). The codon adaptation index (CAI) is an indicator of gene expression, and is normally used to design nucleotide sequences suitable for maximal protein production (Gustafsson et al., 2012). The higher the CAI value the higher the gene expression, and vice versa. The 13 SARS-CoV-2 isolates used in the analysis showed an overall average CAI value of 0.54 ± 0.0003 for the complete coding region, which is comparatively slightly lower than the CAI values of viruses from other studies in relation to *Homo sapiens*, indicating moderate SARS-CoV-2 adaptation to *Homo sapiens* (Butt et al., 2016; Khandia et al., 2019; Kumar et al., 2018). However, at the individual genes level, different CAI values have been observed. The highest CAI value was recorded for the N gene encoding for the nucleocapsid protein (0.62 ± 0.0008) which plays a fundamental role during viral self-assembly (Nelson et al., 2005). ORF7a encoding for the 7a accessory protein recorded the second highest CAI value of 0.58 ± 0.0000, which may play a role in viral assembly or budding events unique to SARS-CoV (Chang et al., 2014). Interestingly, only a small amounts of the envelope protein encoded by the E gene is sufficient to trigger the formation of virus-like particles (Nelson et al., 2005), which may explain its lowest CAI value (0.48 ± 0.001) among other genes.

Relative codon deoptimization index (RCDI) is measured by comparing the codon usage of viruses with that of their hosts. Lower RCDI values indicates higher adaptation of a virus to its host, and vice versa. A study was made on poliovirus in which many synthetic constructs with deoptimized codon usage in their capsid region was developed. Constructs having RCDI values > 2 were not able to survive, unlike other constructs which were able to survive with RCDI values < 2, indicating the poor adaptation of viruses showing higher RCDI values (Mueller et al., 2006). In this study, the average RCDI value for the complete coding region across all isolates was 1.76 ± 0.003, reflecting quite high deoptimization of SARS-CoV-2 codon usage to *Homo sapiens*, which is consistent with CAI results. Moreover, A correlation analysis was performed between the average CAI and RCDI values of SARS-CoV-2 genes revealed a strong negative correlation (r = −0.78, *P* <.05) (Table 1). This result suggests that, an attenuated viral vaccine can be obtained by designing viral genomes with modified codon usage to exhibit higher RCDI and lower CAI values, which will cause a reduction in the rate of SARS-CoV-2 proteins synthesis (Mueller et al., 2006).

## 5. Conclusion

In summary, the findings of this study revealed that the codon usage of SARS-CoV-2 is slightly biased similar to most studied RNA viruses, and the influence of natural selection is more profound than that of mutation pressure in shaping the codon usage pattern of SARS-CoV-2. Other factors including compositional constraints and hydrophobicity also affected SARS-CoV-2 CUB. Overall, the 13 SARS-CoV-2 isolates used in the analysis showed almost identical patterns of codon usage reflecting the small evolutionary changes between them. The findings of this study may help in understanding the underlying factors involved in SARS-CoV-2 evolution, adaptation, and fitness toward *Homo sapiens*.

## Acknowledgement

none.

## Funding

This research did not receive any specific grant from funding agencies in the public, commercial, or no-for-profit sectors.

## Declarations of interest

none.

---

**Supplementary file 1.** Information about the 13 SARS-CoV-2 isolates used in the study [Source: National Centre for Biotechnology Information (NCBI) virus portal; https://www.ncbi.nlm.nih.gov/labs/virus/vssi/#/].

**Supplementary file 2.** Master table containing codon usage bias (CUB) analyses results and nucleotide compositions for each coding sequence in all isolates.

**Supplementary file 3.** Relative synonymous codons usage (RSCU) average values of each codon for the 13 SARS-CoV-2 isolates. Putative preferred or overrepresented codons are highlighted in yellow.

**Supplementary file 4.** The ENc, CAI, RCDI, and tAI average values for each SARS-CoV-2 isolate named by their geo-location (country).

**Supplementary file 5.** Comparison between SARS-CoV-2 and *Homo sapiens* average RSCU values. The most frequently used codon for each amino acid is highlighted in yellow.

## Notes

### Competing Interest Statement

The authors have declared no competing interest.

### Summary of Updates

A section on the materials and methods, as well as on the result were updated, to clarify, without changing any significant results from version 2.

