## Supplementary figures and images for "Insights into The Codon Usage Bias of 13 Severe Acute Respiratory Syndrome Coronavirus 2 (SARS-CoV-2) Isolates from Different Geo-locations"

### Figure S1

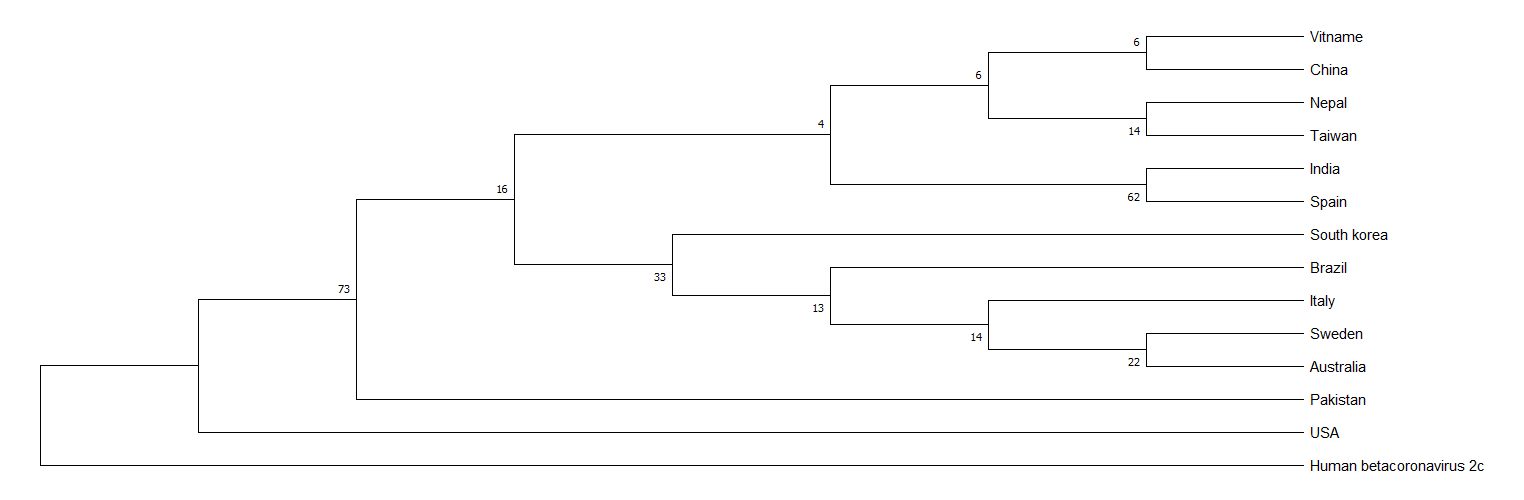
